## Supplementary material for "HIV-1 Vpr combats the PU.1-driven antiviral response in primary human macrophages": All_supplementary

Supplementary Table 1

| **Population** | **Virus treated** | ***gag+/tat+*** | ***gag-/tat-*** |
|---|---|---|---|
| Cluster 0 | 47880 | 14641 | 33239 |
| Cluster 1 | 1539 | 214 | 1325 |
| WT-vpr | 13639 | 6156 | 7483 |
| Vpr-null | 35780 | 8699 | 27081 |

**Supplementary Table 1. MDM infection status by population measured with Gag.** The number of MDMs treated with the indicated virus ($89.6^{wt}$ or $89.6^{\Delta vpr}$) included in our scRNA-seq analysis (Figure 1E). Cells listed as either virus treated, *gag+/tat+* or *gag-/tat-* within each cluster, or within infection exposure (WT = $89.6^{wt}$ and ∆Vpr = $89.6^{\Delta vpr}$) across all three donors.

Supplementary Table 2

| | | | Percent Infected | |
|---|---|---|---|---|
| Donor | Experiment Type | Molecule | 89.6 WT vpr | 89.6 vpr-null |
| 1 | Flow Cytometry | Protein | 65% | 50% |
| 1 | scRNA-seq | mRNA | 71% | 48% |
| 2 | Flow Cytometry | Protein | 45% | 31% |
| 2 | scRNA-seq | mRNA | 53% | 28% |
| 3 | Flow Cytometry | Protein | 52% | 35% |
| 3 | scRNA-seq | mRNA | N/A | (a) 42% |
| 3 | scRNA-seq | mRNA | N/A | (b) 54% |

**Supplementary Table 2. MDM infection rates measured with Gag.** Percent infection of $89.6^{wt}$ and $89.6^{\Delta vpr}$ infected MDMs from each of three donors was determined by quantifying the percent $Gag^{+}$ cells by either flow cytometry (protein) or gene expression levels from scRNA-seq data (mRNA), as indicated by the experiment type and molecule, over the total number of cells analyzed.

### Supplementary Figure 1

##### ETS Family Transcription Factors

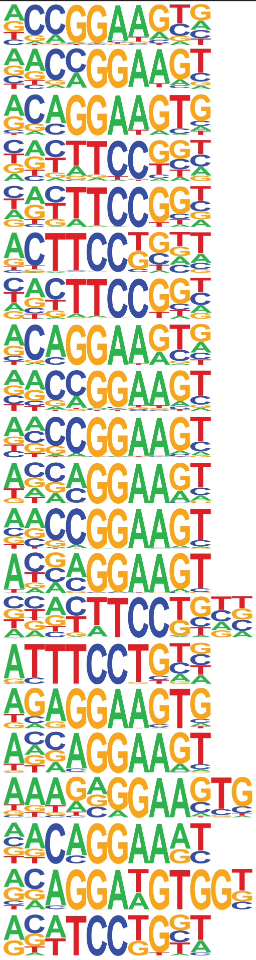

| Motif | TF Name | q-value | # Targets w/ Sequence |
|---|---|---|---|
| | ETV4 | 0.0000 | 1270 |
| | ETV1 | 0.0000 | 1228 |
| | ETS1 | 0.0000 | 987 |
| | Fli1 | 0.0000 | 1229 |
| | Elk4 | 0.0000 | 1038 |
| | Elf4 | 0.0000 | 930 |
| | Elk1 | 0.0000 | 1013 |
| | ERG | 0.0000 | 1135 |
| | GABPA | 0.0000 | 995 |
| | ELF1 | 0.0000 | 925 |
| | EHF | 0.0000 | 876 |
| | ETS | 0.0000 | 674 |
| | ELF5 | 0.0000 | 557 |
| | Etv2 | 0.0000 | 768 |
| | EWS | 0.0001 | 327 |
| | PU.1 | 0.0005 | 316 |
| | ELF3 | 0.0011 | 417 |
| | SpiB | 0.0038 | 180 |
| | EWS:FLI1-fusion | 0.0063 | 534 |
| | ETS:RUNX | 0.0082 | 121 |
| | SPDEF | 0.0121 | 624 |

##### Homeobox Family Transcription Factors

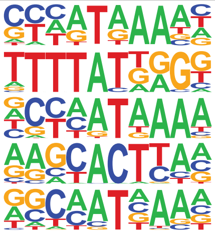

| Motif | TF Name | q-value | # Targets w/ Sequence |
|---|---|---|---|
| | Hoxa13 | 0.0043 | 647 |
| | HOXB13 | 0.0183 | 292 |
| | Hoxd13 | 0.0183 | 1 |
| | Nkx3.1 | 0.0255 | 746 |
| | Hoxa9 | 0.0355 | 796 |

##### bZIP Family Transcription Factors

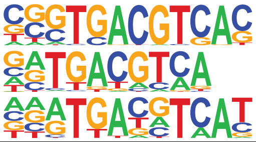

| Motif | TF Name | q-value | # Targets w/ Sequence |
|---|---|---|---|
| | CRE | 0.0000 | 385 |
| | Atf1 | 0.0043 | 434 |
| | CREB5 | 0.0632 | 203 |

##### NRF Family Transcription Factors

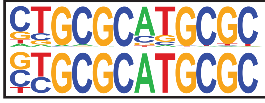

| Motif | TF Name | q-value | # Targets w/ Sequence |
|---|---|---|---|
| | NRF1 | 0.0000 | 544 |
| | NRF | 0.0003 | 1 |

##### CCAAT Family Transcription Factors

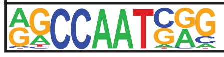

| Motif | TF Name | q-value | # Targets w/ Sequence |
|---|---|---|---|
| | NFY | 0.0000 | 782 |

##### ETS/IRF Family Transcription Factors

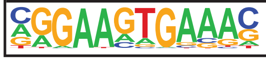

| Motif | TF Name | q-value | # Targets w/ Sequence |
|---|---|---|---|
| | PU.1-IRF | 0.0004 | 670 |

##### IRF Family Transcription Factors

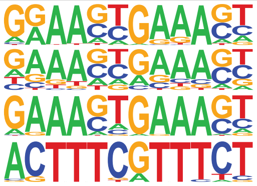

| Motif | TF Name | q-value | # Targets w/ Sequence |
|---|---|---|---|
| | IRF8 | 0.0006 | 209 |
| | IRF2 | 0.0049 | 75 |
| | IRF1 | 0.0453 | 68 |
| | T1ISRE | 0.0504 | 8 |

##### ZF Transcription Factors

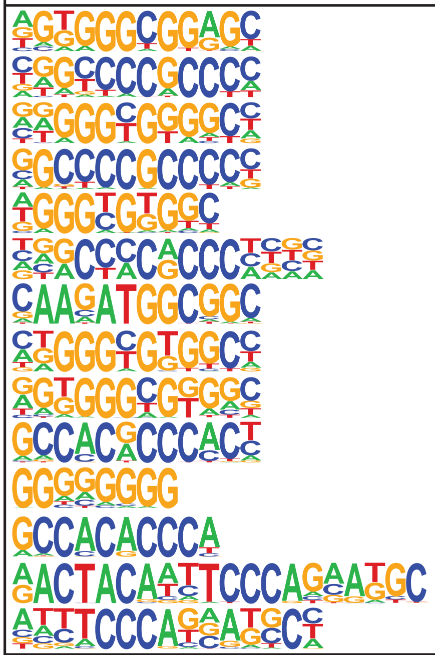

| Motif | TF Name | q-value | # Targets w/ Sequence |
|---|---|---|---|
| | Sp5 | 0.0000 | 1786 |
| | Sp2 | 0.0000 | 2047 |
| | KLF1 | 0.0000 | 1607 |
| | Sp1 | 0.0000 | 1125 |
| | KLF5 | 0.0000 | 1787 |
| | KLF3 | 0.0000 | 1089 |
| | YY1 | 0.0000 | 203 |
| | KLF6 | 0.0000 | 1497 |
| | KLF14 | 0.0000 | 2044 |
| | Klf9 | 0.0001 | 780 |
| | Maz | 0.0012 | 1644 |
| | Klf4 | 0.0056 | 531 |
| | GFY | 0.0183 | 200 |
| | ZNF143\|ST | 0.0567 | 275 |

##### bHLH Family Transcription Factors

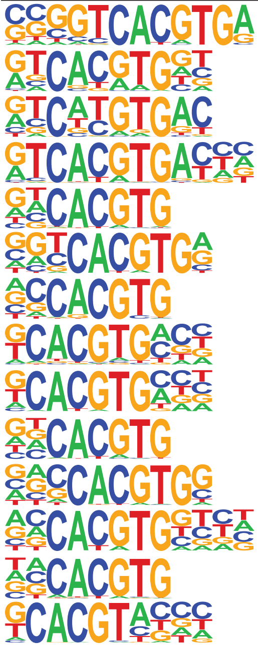

| Motif | TF Name | q-value | # Targets w/ Sequence |
|---|---|---|---|
| | E-box | 0.0000 | 212 |
| | Usf2 | 0.0000 | 290 |
| | MITF | 0.0000 | 505 |
| | TFE3 | 0.0000 | 160 |
| | CLOCK | 0.0000 | 402 |
| | USF1 | 0.0002 | 371 |
| | c-Myc | 0.0006 | 495 |
| | bHLHE41 | 0.0036 | 888 |
| | bHLHE40 | 0.0041 | 280 |
| | BMAL1 | 0.0043 | 679 |
| | n-Myc | 0.0150 | 432 |
| | Max | 0.0228 | 347 |
| | NPAS | 0.0254 | 612 |
| | HIF2a | 0.0636 | 246 |

##### THAP Family Transcription Factors

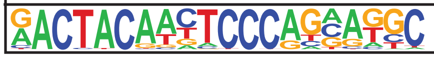

| Motif | TF Name | q-value | # Targets w/ Sequence |
|---|---|---|---|
| | Ronin | 0.0227 | 185 |

**Supplementary Figure 1. Transcription factors and their predicted binding motifs in downregulated genes in MDMs expressing Vpr.** Motifs, transcription factor names, significance (FDR q-value), and number of target genes identified by HOMER analysis of genes downregulated in the presence of Vpr (Figure 1E) from several transcription factor families.

Supplementary Figure 2

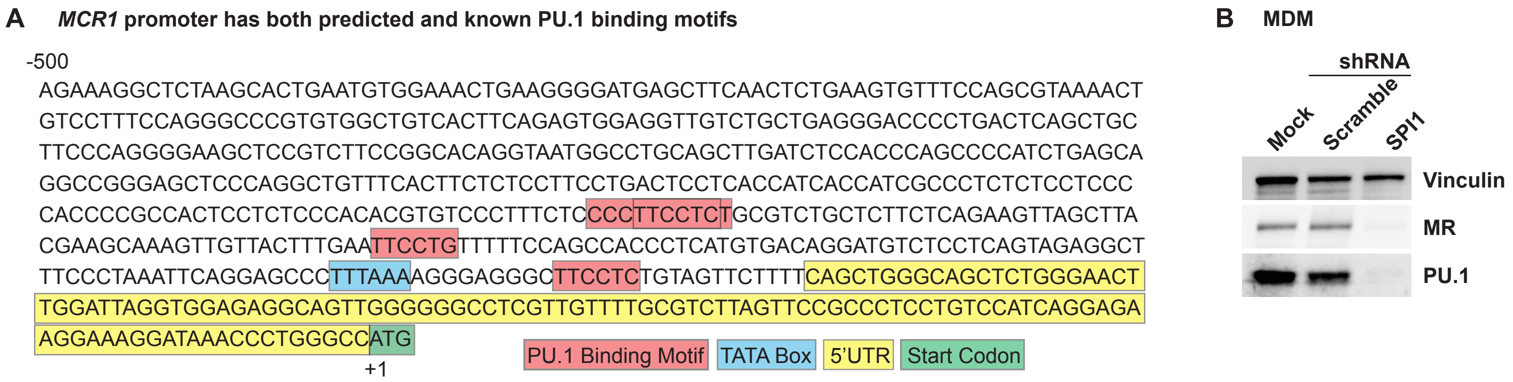


**Supplementary Figure 2. The MRC1 promoter contains multiple PU.1 binding motifs.** (A) The first 500 bp upstream of the start codon in *MRC1*, the gene that codes for mannose receptor. Previously reported PU.1 binding motifs are outlined in red. The double-box PU.1 motif was identified through inputting the HOMER generated PU.1 motif parameters into FIMO (Find Individual Motif Occurrences). The TATA box is outlined in blue, the 5'UTR in yellow, and the start codon in green. (B) Immunoblot analysis from MDMs stably expressing the indicated shRNAs, n = 2.

Supplementary Figure 3

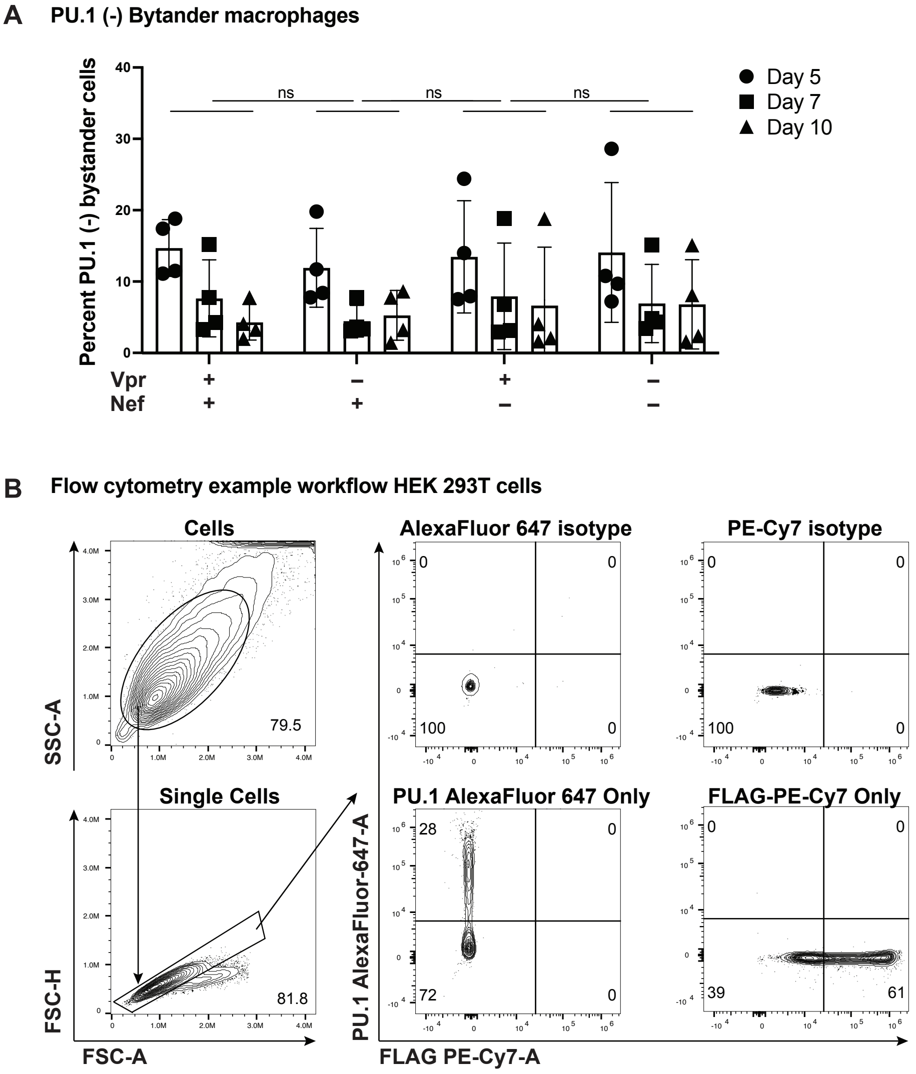


**Supplementary Figure 3. PU.1 is not significantly reduced in bystander MDMs at 5-, 7-, and 10-days post infection with replication defective (non-spreading) HIV constructs.** (A) Summary graph showing the percentage of infected ($GFP^-$) cells that do not express PU.1 as determined by flow cytometry as depicted in Figure 4B. The mean +/- standard deviation from n=4 independent donors is shown for each time point. P values were determined using an analysis of variance (ANOVA) with Tukey’s multiple comparisons test; ns = not significant. (B) Gating strategy used for flow cytometry in Figures 4 and 5, and Supplementary Figure 4.

### Supplementary Figure 4

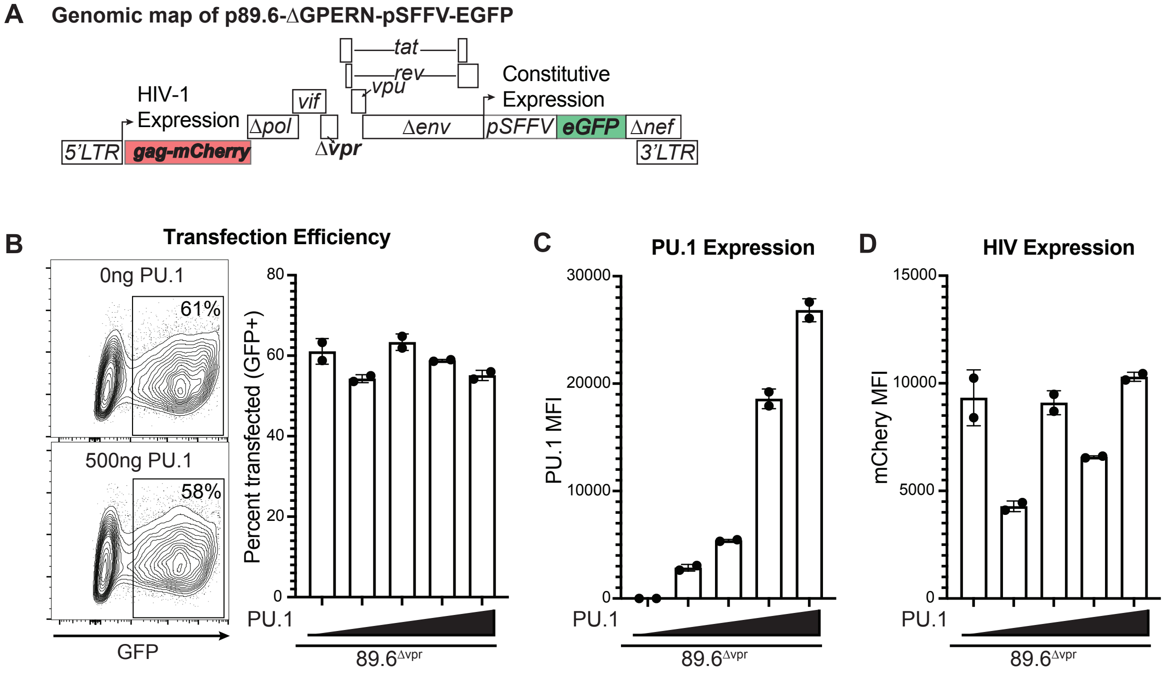


**Supplementary Figure 4. PU.1 does not alter HIV-LTR activity.** (A) Genomic map for 89.6-derived HIV-1 fluorescent reporter virus (89.6-ΔGPERN-pSFFV-EGFP). (B) Representative flow plots and bar graph assessing transfection rate via GFP expression in HEK 293T cells transfected with the indicated amount of PU.1 construct from Figure 2E plus 500ng of 89.6-ΔGPERN-pSFFV-EGFP from (A). (C) Summary graph of PU.1 expression in GFP+ cells from (B). PU.1 levels were assessed using intracellular staining as described in Methods and measured by flow cytometry. (D) Summary graph of HIV-LTR activity as assessed by mCherry MFI from the same cells as in (C), measured flow cytometrically. All conditions were transfected with 500ng of HIV expression plasmid plus 0, 1, 10, 100, or 500ng of PU.1 plasmid. MFI = mean fluorescence intensity.
